# Distinct stages in the recognition, sorting and packaging of proTGFα into COPII coated transport vesicles

**DOI:** 10.1101/040410

**Authors:** Pengcheng Zhang, Randy Schekman

## Abstract

In addition to its role in forming vesicles from the endoplasmic reticulum (ER), the coat protein complex II (COPII) is also responsible for selecting specific cargo proteins to be packaged into COPII transport vesicles. Comparison of COPII vesicle formation in mammalian systems and in yeast suggested that the former employs more elaborate mechanisms for cargo recognition, presumably to cope with a significantly expanded repertoire of cargo that transits the secretory pathway. Using proTGFα, the transmembrane precursor of transforming growth factor alpha (TGFα), as a model cargo protein, we demonstrate in cell-free assays that at least one auxiliary cytosolic factor is specifically required for the efficient packaging of proTGFα into COPII vesicles. Using a knockout HeLa cell line generated by CRISPR/Cas9, we provide functional evidence showing that a transmembrane protein, Cornichon-1 (CNIH), acts as a cargo receptor of proTGFα. We show that both CNIH and the auxiliary cytosolic factor(s) are required for efficient recruitment of proTGFα to the COPII coat *in vitro*. Moreover, we provide evidence that the recruitment of cargo protein by the COPII coat precedes and may be distinct from subsequent cargo packaging into COPII vesicles.

**Abbreviations:**

CNIH: Cornichon

## Introduction

The endoplasmic reticulum (ER) is the entry point for newly synthesized proteins into the secretory pathway, and the coat protein complex II (COPII) is the membrane coat that is responsible for recruiting and transporting cargo from the ER via COPII vesicles. The COPII coat consists of five core components: The small GTPase Sar1, the Sec23/24 heterodimer, and the Sec13/31 heterotetramer. Cytosolic Sar1 is activated when it exchanges bound GDP for GTP, a process mediated by the guanine nucleotide exchange factor (GEF) Sec12, which is an ER resident protein (Nakano *et al*., 1988; Barlowe and Schekman, 1993). Activated Sar1 extends an amphipathic N-terminal a-helix and associates with the ER membrane, to which it recruits the Sec23/24 heterodimer, forming the pre-budding complex (Kuehn *et al*., 1998; Huang *et al*., 2001). After assembly of the pre-budding complex, the Sec13/31 heterotetramer is recruited to the ER where it polymerizes to guide membrane curvature and vesicle formation (Antonny *et al*., 2001).

The selection and enrichment of COPII cargo proteins is achieved by interaction with the COPII coat, mainly via the Sec24 subunit (Miller *et al*., 2002, 2003). In principle, transmembrane cargo proteins can interact directly with Sec24 via their cytoplasmic domains. Indeed this has been observed for a number of well characterized COPII cargo proteins (Mossessova *et al*., 2003; Wendeler *et al*., 2007). However, in the course of investigating the molecular requirements for COPII packaging of certain mammalian transmembrane proteins, we detected the involvement of one or more cytosolic factors in addition to the COPII coat.

Transforming growth factor alpha (TGFα) was identified as a secreted peptide that had cell transforming properties. TGFα is synthesized as a small type I transmembrane protein with a short cytoplasmic C-terminus that has a PDZ binding motif (PBM) at the C-terminus. The N-terminal luminal domain contains glycosylation sites and an EGF-like domain. At the cell surface, proTGFα undergoes two sequential proteolytic cleavages that results in the release of the soluble TGFα ligand into the extracellular space (Singh and Coffey, 2014). In HeLa cells, proTGFα was reported to interact in the ER with Cornichon-1 (hereafter referred to as Cornichon and abbreviated as CNIH for simplicity). A luminal loop of CNIH was shown to interact with the luminal EGF-like domain of proTGFα in the ER (Castro *et al*., 2007). CNIH was proposed to act as a cargo receptor for proTGFα, but overexpression of CNIH-HA paradoxically led to more retention of proTGFα in the ER, making it unclear whether CNIH plays a direct role in the ER export of proTGFα *in vivo* (Castro *et al*., 2007). In addition, mutation of the cytoplasmic C-terminal valine to glycine (V160G) was shown in multiple studies to significantly impede the ER export kinetics of proTGFα, suggesting a cytoplasmic component involved in proTGFα trafficking (Bosenberg *et al*., 1992; Fernández-Larrea *et al*., 1999; Ureña *et al*., 1999). Using a cell-free reaction that reproduces the cargo-selective packaging of mammalian membrane cargo proteins into COPII vesicles, we probed the requirement for auxiliary cytosolic factors in the ER export of specific mammalian cargo, as well as the function of CNIH in mammalian cells.

## Results

### Evidence for Existence of Auxiliary Cytosolic Factors

Cytosol prepared from fresh rat livers was sufficient to support packaging of both pro/HA-TGFα and ERGIC53, a lectin that cycles between the ER and ER-Golgi intermediate compartment (ERGIC), into COPII vesicles made in a cell-free vesicle budding reaction. The control protein ribophorin I, an ER resident protein and component of the eukaryotic oligosaccharide transferase complex, was absent in the vesicle fractions, indicating the preservation of ER membrane integrity (Figure 1A, lanes 3 & 4.). Cytosol prepared from frozen rat livers (livers were flash-frozen in liquid nitrogen and then thawed slowly at 4°C), in contrast, was inactive in the budding reaction (Figure 1A, lanes 5 & 6). Purified COPII components supported the efficient packaging of ERGIC53, but not pro/HA-TGFα (Figure 1A, lane 7), suggesting the requirement of a cargo-selective cytosolic factor(s) for the packaging of the latter. Addition of freeze-thaw inactivated cytosol and purified COPII components restored the packaging of pro/HA-TGFα into COPII vesicles (Figure 1A, lanes 8-11). Thus, we speculated that the freeze-thaw treated cytosol may have lost COPII activity but retained an additional factor essential for the packaging of proTGFα.

**Figure 1.**
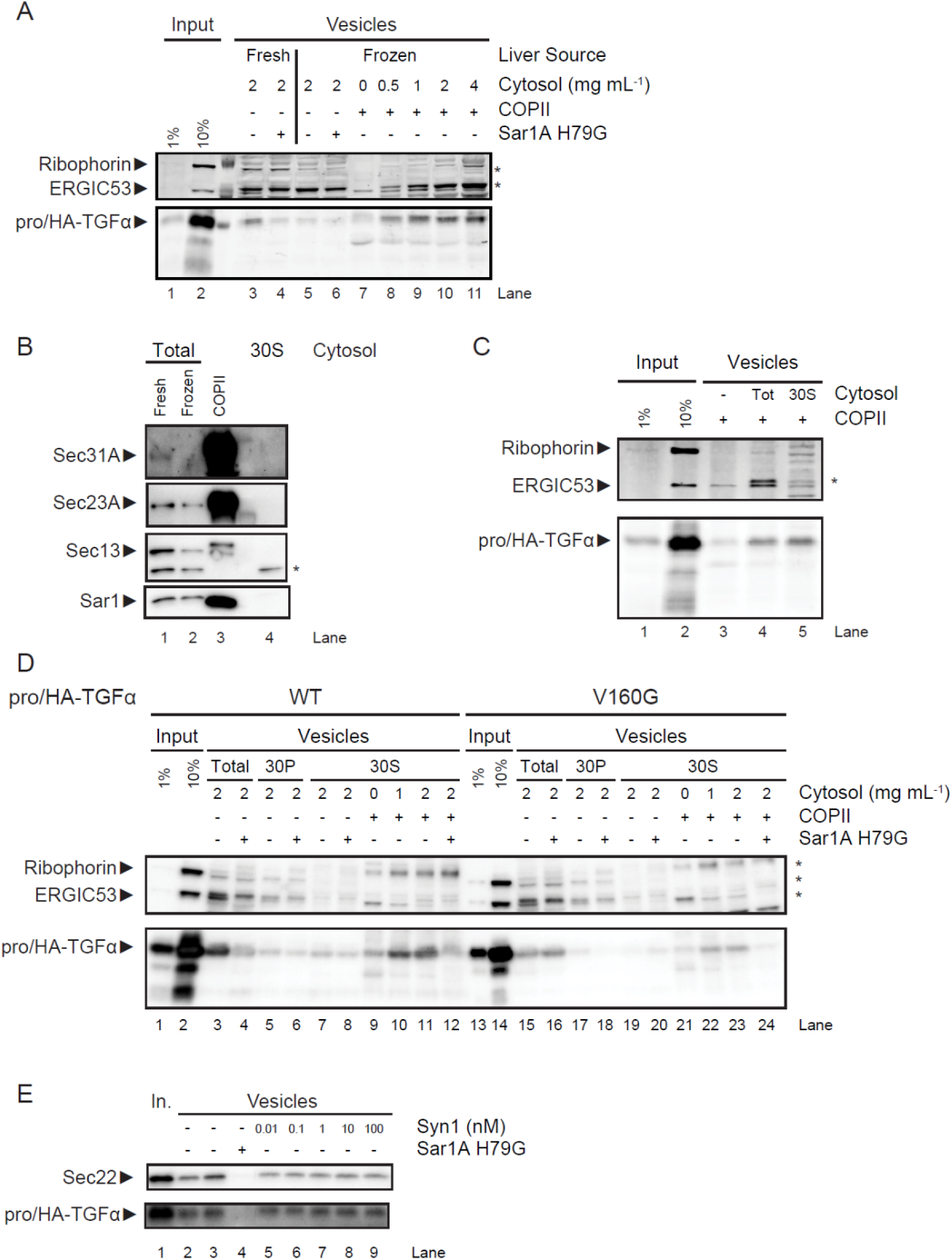
ProTGFα requires an auxiliary cytosolic factor for efficient COPII packaging. HeLa cells were transfected with plasmids expressing pro/HA-TGFα and membranes were harvested the next day for use in *in vitro* budding reactions. (A) LiCor scan of *in vitro* budding reaction. Rat liver cytosol was prepared from fresh or frozen rat livers as indicated. Sar1A H79G is a GTPase mutant that is a dominant negative inhibitor of *in vitro* budding. (B) Immunoblot of various cytosol fractions and purified COPII samples. (C) LiCor scan of *in vitro* budding reaction. (D and E) Immunoblots of *in vitro* budding reactions. Tot: Total cytosol extract. 30P: Pellet fraction of cytosol after 30% ammonium sulfate precipitation. 30S: Supernatant fraction of cytosol after 30% ammonium sulfate precipitation. Syn1: Purified recombinant human Syntenin-1. Asterisks: Unrelated bands.

Although cytosol prepared from frozen rat livers displayed minimal budding activity, it still contained significant amounts of the COPII subunits (Figure 1B, lanes 1 & 2). To rule out the possibility that the pro/HA-TGFα-specific packaging activity was due to the residual COPII components, proteins from the inactive cytosol were mixed with 30% ammonium sulfate under conditions where COPII proteins are precipitated (Kim *et al*., 2005), which reduced COPII levels below the detection limit (Figure 1B, lane 4). Vesicle budding reactions supplemented with a dialyzed aliquot of the 30% supernatant fraction (30S) showed specific enhancement of pro/HATGFα packaging (Figure 1C, compare lanes 3 & 5), indicating that the pro/HA-TGFα-specific activity was not attributable to known COPII components, but possibly to at least one auxiliary cytosolic factor specifically required in the packaging of pro/HA-TGFα. Notably, while ERGIC53 packaging was enhanced by total cytosol extract, the 30S cytosol fraction did not have an effect in this regard (Figure 1C, lanes 4 & 5).

Further experiments were conducted to confirm that the observed pro/HA-TGFα packaging activity was a COPII-dependent process. The 30S cytosol fraction was unable to support packaging of either ERGIC53 or pro/HA-TGFα in the absence of purified COPII (Figure 1D, lane 7). Supplementing purified COPII components with the 30S fraction greatly enhanced pro/HA-TGFα packaging efficiency (Figure 1D, lanes 9-11), and this activity was inhibited when SarlA H79G, a dominant negative inhibitor of COPII budding *in vitro*, was included in the reaction (Figure 1D, lane 12), thus confirming that the pro/HA-TGFα-specific activity of the 30S fraction was dependent on COPII.

To test the physiological relevance of our *in vitro* observations, we examined the behavior of the V160G mutant form of proTGFα, which exhibited trafficking defects (Briley *et al*., 1997). When tested in the *in vitro* budding assay, the pro/HA-TGFα V160G was packaged inefficiently into vesicles (Figure 1D, compare lane 15 and lanes 3 & 16). And as expected, supplementing purified COPII with the 30S cytosolic fraction resulted in inefficient packaging of pro/HA-TGFα V160G as well (Figure 1D, compare lanes 22 & 23 and lanes 10 & 11), thus the mutant behaved *in vitro* as it did *in vivo*.

The PDZ domain containing protein Syntenin-1 (Syn1) was previously identified as an interaction partner of proTGFα. Syn1 interacted with the wild type C-terminus of proTGFα, but not with the V160G mutant form. Hence it was proposed to function in the early steps of proTGFα trafficking (Fernández-Larrea *et al*., 1999). In light of the observations above, we tested whether Syn1 acted as the auxiliary cytosolic factor. Purified recombinant Syn1 was supplemented to *in vitro* budding reactions, but no significant effect on pro/HA-TGFα budding was observed (Figure 1E). Thus our *in vitro* evidence does not support a role for Syn1 as the auxiliary factor. However, it is possible that the purified Syn1 is inactive, and our results do not rule out the possibility that Syn 1 might act in coordination with other cytosolic factors.

### Cornichon functions as a cargo receptor for proTGFα

In *Drosophila*, Cornichon has been implicated in the ER export of Gurken (Roth *et al*., 1995; Bökel *et al*., 2006), a single-pass transmembrane protein that also contains an extracellular EGF-like domain. The Cornichon homolog in yeast, Erv14p, has been characterized to function as a cargo receptor for the ER export of Axl2p (Powers, 1998; Powers and Barlowe, 2002), among other client proteins (Nakanishi *et al*., 2007; Herzig *et al*., 2012).

CNIH has been proposed to perform a similar function in mammalian cells, but previous characterization of its role in trafficking *in vivo* was inconclusive (Castro *et al*., 2007). To ask whether CNIH might indeed play such a role, we generated a CNIH knockout (KO) cell line in HeLa cells using CRISPR/Cas9. Seven clones were expanded and analyzed for expression of CNIH (Figure 2A). Genomic sequencing confirmed that clone #3 was a bona fide genomic knockout of CNIH, and this cell line was used in subsequent experiments.

**Figure 2.**
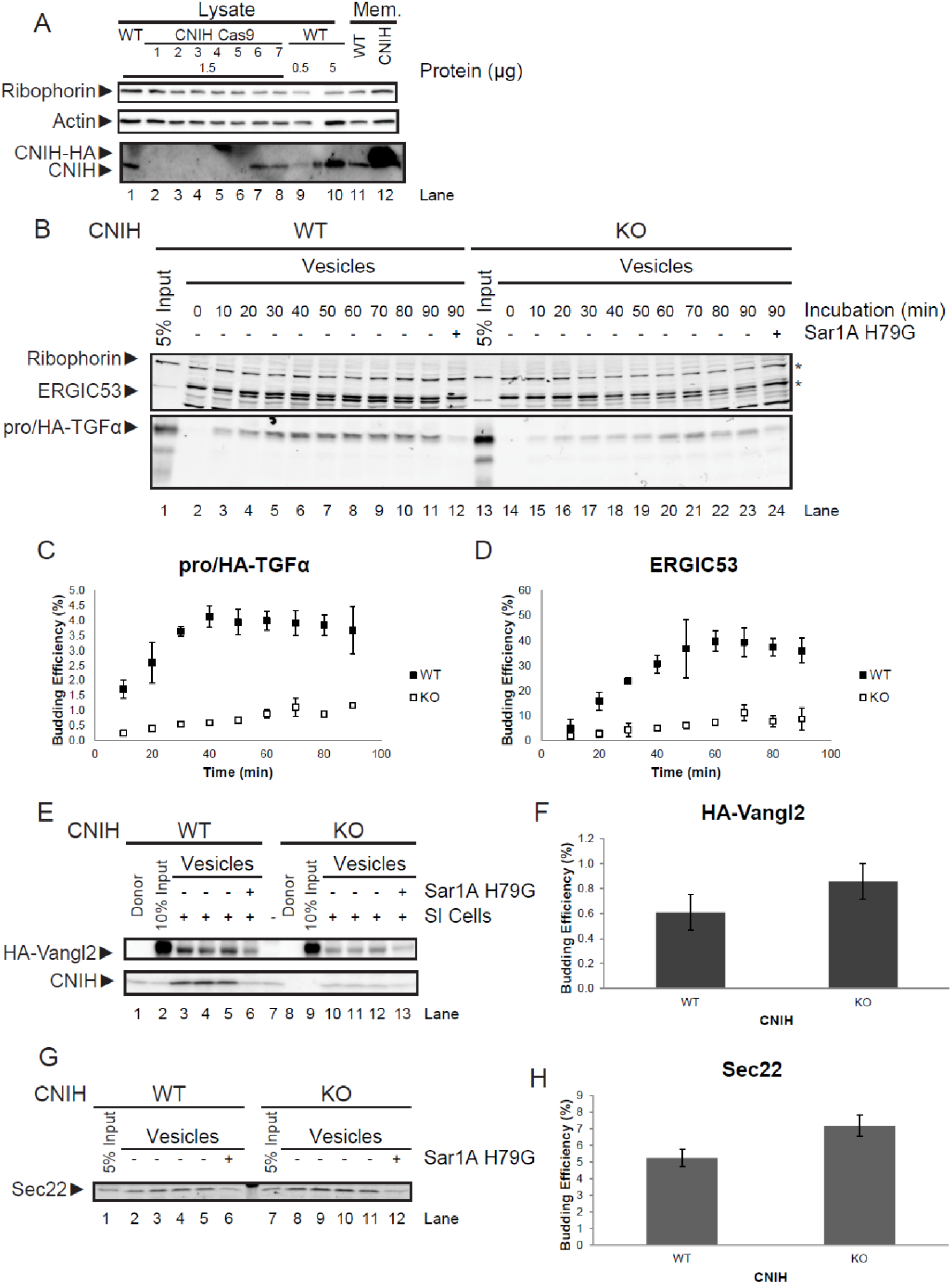
Loss of CNIH impairs ER export of proTGFα. (A) Immunoblot of HeLa cell samples. Lysate: Total cell lysate. WT: Wild type. CNIH Cas9: Cells that have been treated for modification at the CNIH locus by CRISPR/Cas9. Mem.: Permeabilized wild type cells that were either untreated (WT) or transfected with a plasmid encoding CNIH-HA (CNIH). (B) LiCor scan of an *in vitro* budding reaction. (C and D) Quantification of budding efficiency of pro/HA-TGFα (C) and ERGIC53 (D). (E) Immunoblot of *in vitro* budding reactions. Donor: Permeabilized HeLa cells that were not subjected to *in vitro* translation. (G) LiCor scan of *in vitro* budding reactions. (F and H) Quantification of budding efficiency of HA-Vangl2 (F) and Sec22 (H). Quantifications are represented as mean±SEM (p>0.05 in both cases). Asterisks: Unrelated bands.

We first assessed the effect of CNIH KO on proTGFα trafficking. CNIH KO resulted in diminished budding efficiency of pro/HA-TGFα, but also resulted in diminished ERGIC53 budding efficiency (Figure 2, B–D). However, CNIH KO did not significantly affect the trafficking of Sec22, a v-SNARE which cycles between the ER and ERGIC, and Vangl2, a transmembrane protein involved in cell signaling in the planar cell polarity pathway (Figure 2, E–H), suggesting that CNIH KO did not result in general disruption of COPII vesicle formation. Importantly, CNIH itself was also packaged very efficiently into COPII vesicles (Figure 2E). Together these data point to a role for CNIH in the ER export of proTGFα.

### Reintroducing Cornichon into CNIH KO HeLa cells partially rescues proTGFα budding phenotype

To confirm that the proTGFα budding defects observed in CNIH KO membranes could be attributed to the absence of CNIH, rescue experiments were performed. Full-length human CNIH was put under control of a tetracycline inducible promoter and stably integrated into CNIH KO HeLa cells. A moderate level of CNIH expression was induced by addition of doxycycline (Figure 3E). Cells were then processed for use in the *in vitro* budding reactions.

In membranes prepared from wild type HeLa cells, pro/HA-TGFα was efficiently exported from the ER, as was CNIH (Figure 3A). In membranes prepared from CNIH KO cells, packaging of pro/HA-TGFα into COPII vesicles was significantly impaired (Figure 3C). When a moderate level of CNIH expression was reintroduced into CNIH KO cells (rescue), pro/HATGFα budding efficiency was significantly increased above KO levels, although not restored to WT levels (Figure 3, B and D).

**Figure 3.**
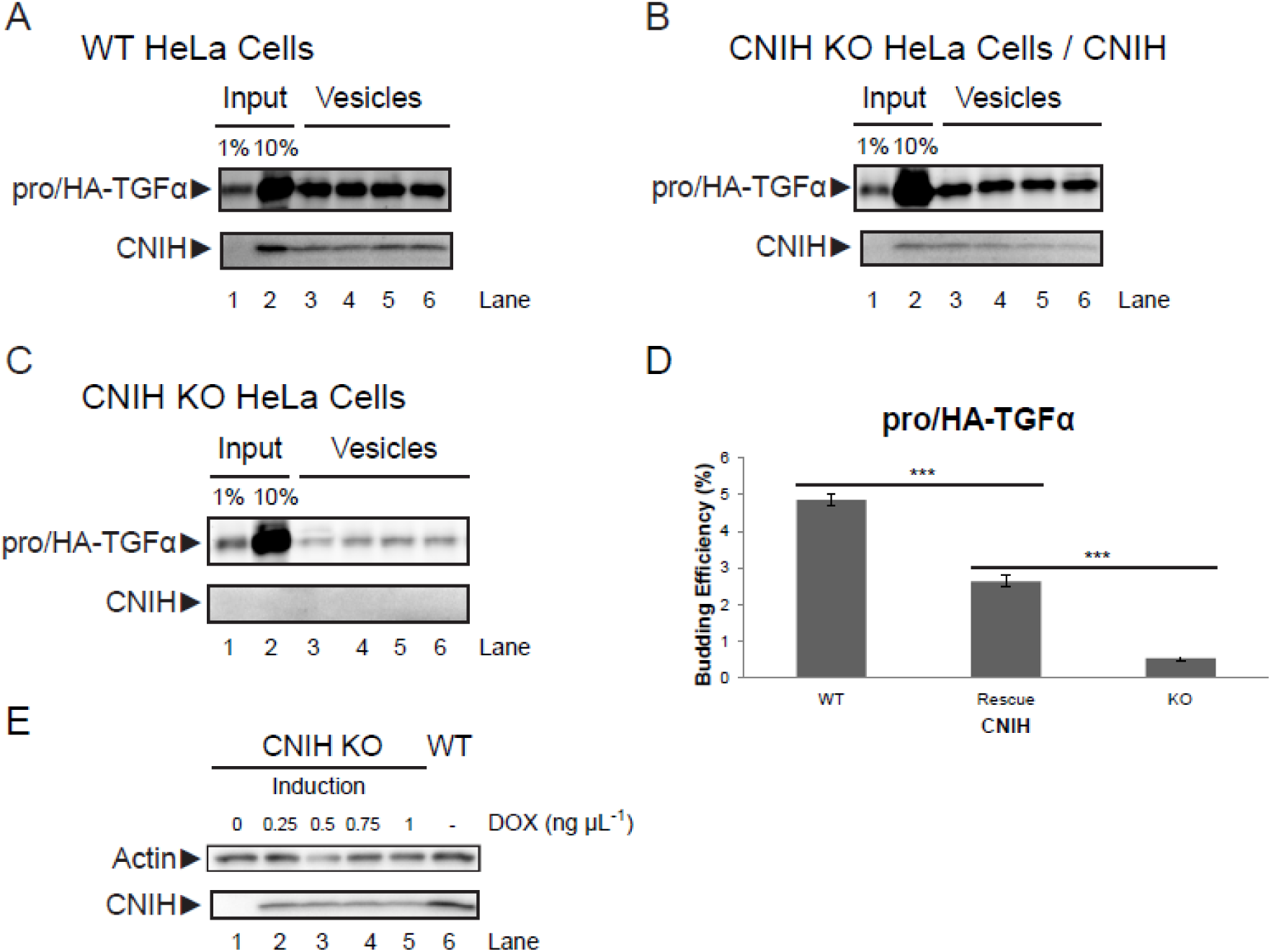
Expression of CNIH in CNIH KO HeLa cells partially rescues proTGFα budding phenotype. (A-C) *In vitro* budding reactions. CNIH was detected using immunoblotting whereas pro/HA-TGFα was detected using LiCor. Membranes were prepared from: wild type HeLa cells expressing pro/HA-TGFα (A), CNIH KO HeLa cells expressing pro/HA-TGFα & CNIH induced by 1 ng μl^−1^ doxycycline (B), or CNIH KO HeLa cells expressing pro/HA-TGFα without doxycycline induction (C). Lanes 3-6 in panels A, B, & C are quadruplicate budding reactions, respectively. (D) Quantification of proTGFα budding efficiency. Results are represented as mean±SEM (***: p<0.001). (E) Immunoblot showing the induction of CNIH expression. Cells were treated with indicated amounts of doxycycline for 24h, and then lysates were harvested for analysis. DOX: doxycycline.

The difference in proTGFα budding efficiency in wild type and rescue membranes may be attributed to the lower CNIH expression levels in the rescue cells. Higher expression levels of CNIH led to ER retention (Figure S1), thus making it challenging to approximate wild type levels of CNIH expression in CNIH KO cells. Despite this drawback, the efficiency of proTGFα budding showed a clear dependence on CNIH levels in the donor membranes (Fig. 3D).

### Cytosol prepared from CNIH KO HeLa cells showed enhanced COPII activity

In the *in vitro* budding reactions, pro/HA-TGFα budding efficiency was significantly reduced in membranes prepared from CNIH KO HeLa cells (Figures 2 and 3). However, immunofluorescence microscopy showed no significant difference in steady state proTGFα localization in WT & CNIH KO cells. In addition, no significant difference in proTGFα trafficking kinetics was observed between WT & CNIH KO cells (Figure S2).

Notably, the experiments in Figures 2 and 3 were done using HeLa cells as membrane source and rat liver cytosol as COPII source. The observations made *in vitro* and *in vivo* suggested the possibility of some compensatory effect attributable to the cytosolic protein content of CNIH KO HeLa cells as an explanation of this discrepancy.

To test this idea, cytosol was prepared from WT and CNIH KO HeLa cells and used in the *in vitro* budding reaction. As shown in Figure 4, cytosol prepared from CNIH KO cells displayed significantly higher COPII activity than that prepared from WT cells (Figure 4A). When incubated with CNIH KO membranes, cytosol prepared from CNIH KO cells significantly increased the budding efficiency of pro/HA-TGFα compared to WT cytosol (Figure 4B). Sec22 budding was also enhanced but to a lesser degree (Figure 4C).

**Figure 4.**
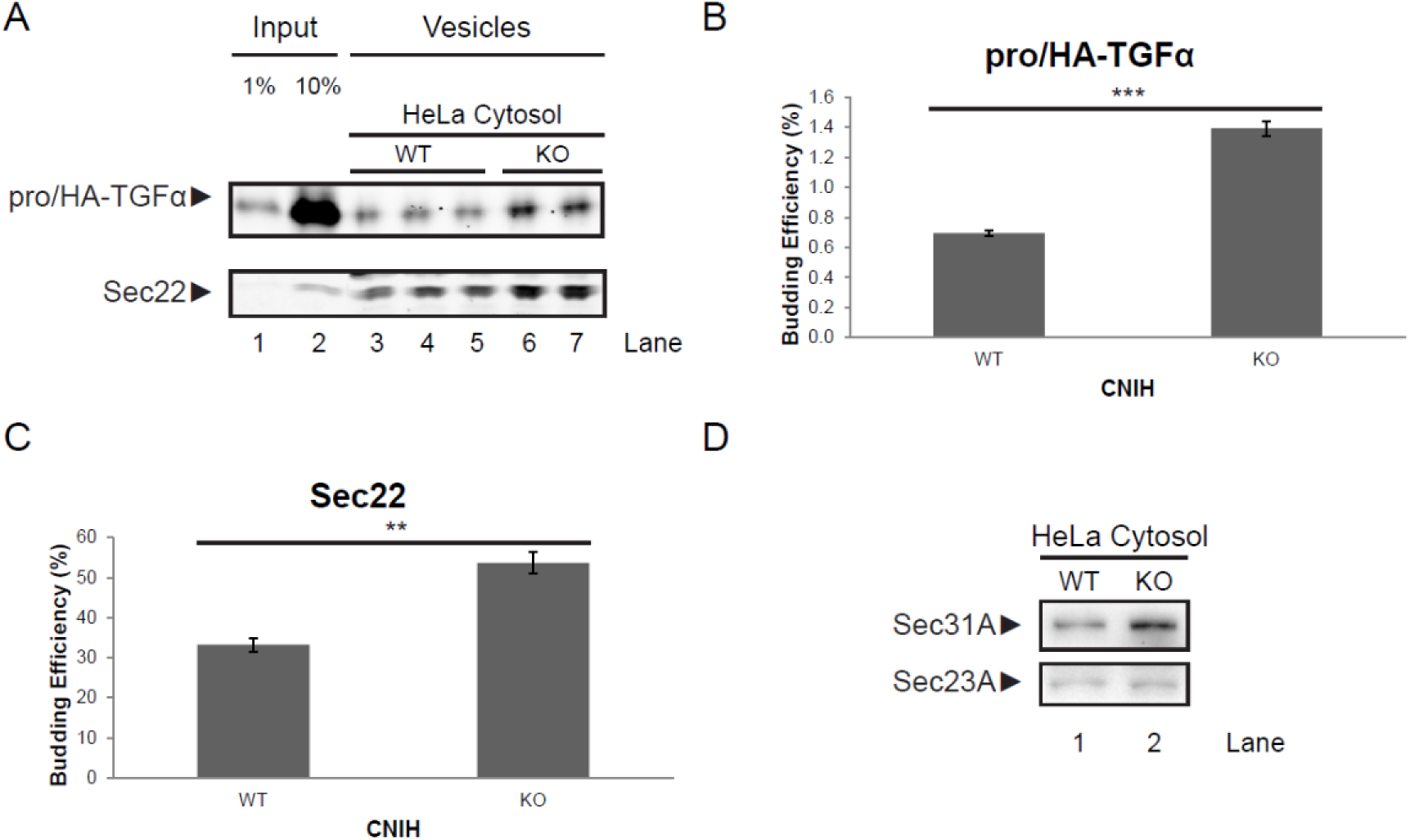
Cytosol from CNIH KO HeLa cells displays higher COPII activity *in vitro*. (A) LiCor scan of an *in vitro* budding reaction. WT HeLa cells were transiently transfected with plasmids encoding pro/HA-TGFα, and membranes were harvested for use in the *in vitro* budding reaction. Cytosol was harvested either from WT HeLa cells or CNIH KO HeLa cells and used in the *in vitro* budding reaction. Lanes 3-5 are triplicates of budding reactions done using cytosol prepared from WT HeLa cells. Lanes 6-7 are duplicates of budding reactions done using cytosol prepared from CNIH KO HeLa cells. (B and C) Quantification of budding efficiency of pro/HA-TGFα (B) and Sec22 (C). Quantifications are represented as mean±SEM (***: p<0.001; **: p<0.01). (D) Immunoblot of cytosol prepared from wild type (WT) or CNIH KO (KO) HeLa cells.

Analysis of cytosolic protein content showed that the expression of Sec31A was increased in CNIH KO cells compared to WT, whereas Sec23A levels remained similar (Figure 4D). This observation also supports the idea that CNIH KO cells may have undergone adaptation in response to the loss of CNIH.

### proTGFα recruitment to pre-budding complex requires cooperation of cytosolic factor(s) and CNIH

The apparent involvement of both a transmembrane cargo receptor and a cytosolic factor(s) in the ER export of proTGFα prompted us to investigate the mechanism of proTGFα recruitment to the pre-budding complex using the COPII recruitment assay, which allowed us to isolate the pre-budding complex and analyze its contents.

To confirm that the trafficking defect of pro/HA-TGFα V160G was not a result of defective binding to CNIH, coimmunoprecipitation experiments were performed and demonstrated that pro/HA-TGFα V160G bound to CNIH as efficiently as wild type. The interaction between CNIH and pro/HA-TGFα was specific, as no interaction was detected between CNIH and HA-Vangl2 (Figure 5A, lanes 7-9. In lanes 7 & 8, an unrelated band appears at an apparent mobility similar to HA-Vangl2.).

**Figure 5.**
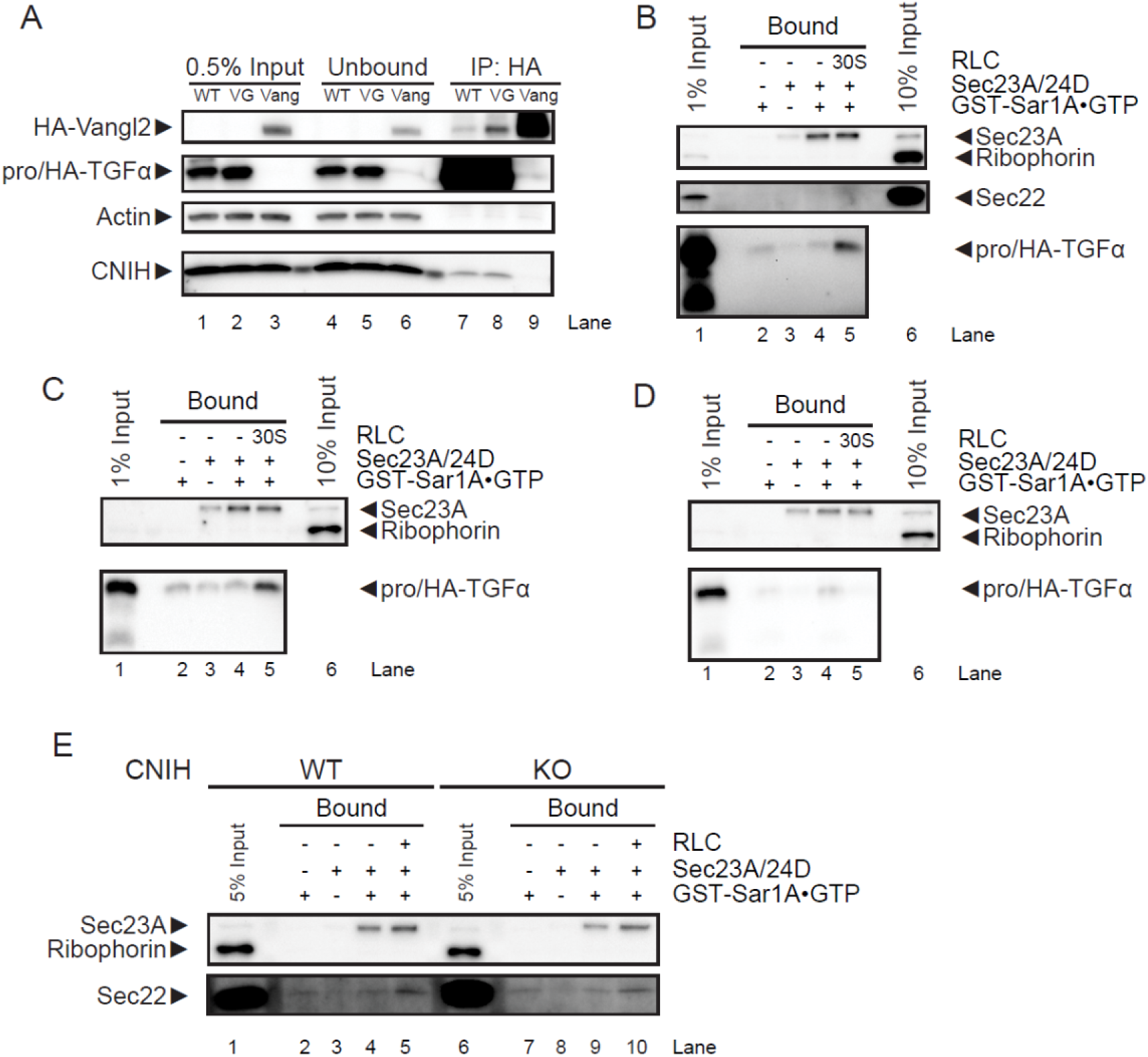
CNIH and cytosolic factor(s) required for proTGFα recruitment to the pre-budding complex. (A) HeLa cells were transiently transfected with plasmids expressing pro/HA-TGFα wild type (WT), V160G mutant (VG), or HA-Vangl2 (Vang) as indicated, and immunoprecipitation was performed using anti-HA antibodies. (B-E) Immunoblots of COPII recruitment assays. Recruitment of pro/HA-TGFα was examined using membranes prepared from wild type HeLa cells expressing proTGFα WT (B) or V160G (C), or using membranes prepared from CNIH KO HeLa cells expressing proTGFα WT (D). Recruitment of Sec22 was compared in HeLa wild type (WT) membranes or CNIH KO (KO) membranes (E).

HeLa cells were transiently transfected with plasmids encoding pro/HA-TGFα WT and membranes were harvested the next day for use in the COPII recruitment assay. The pre-budding complex was captured using GST-Sar1A-GTP-restricted mutant protein, and isolated by GST pulldown (For details, see Materials and Methods). When the pre-budding complex was assembled using purified Sec23A/24D heterodimer, pro/HA-TGFα was not recruited to the pre-budding complex (Figure 5B, lane 4). Supplementing the 30% supernatant fraction (30S) to the reaction resulted in recruitment of pro/HA-TGFα WT to the pre-budding complex, demonstrating the requirement of cytosolic factor(s) in this process (Figure 5B, lane 5). Notably, while pro/HATGFα was recruited to the pre-budding complex, Sec22 was not, indicating cargo specificity. HeLa cells were then transiently transfected with plasmids encoding pro/HA-TGFα V160G, and processed for use in the COPII recruitment assay. Interestingly, although the V160G mutant was defective for packaging into COPII vesicles, it was still recruited to the pre-budding complex in the presence of purified Sec23A/24D and the 30S fraction (Figure 5C, lane 5). Finally, HeLa CNIH KO cells were transiently transfected with plasmids encoding pro/HA-TGFα WT and processed for use in the COPII recruitment assay. Removal of CNIH from HeLa membranes prevented pro/HA-TGFα WT from being recruited to the pre-budding complex (Figure 5D, lane 5). As a control, the impact of CNIH KO on Sec22 recruitment was analyzed. Whereas supplementing the 30S fraction to purified Sec23A/24D did not support Sec22 recruitment to the pre-budding complex, supplementing total rat liver cytosol did (Figure 5E, lane 5). This can be explained by the fact that Sec22 specifically interacts with isoforms A & B of Sec24 (Mancias and Goldberg, 2008). Total rat liver cytosol contained all four Sec24 isoforms, and thus was able to support Sec22 recruitment to the pre-budding complex. In contrast, the 30S fraction did not contain Sec24, and thus Sec22 was not recruited to the pre-budding complex. More importantly, CNIH KO did not affect Sec22 recruitment to the pre-budding complex, supporting a specific role for CNIH in proTGFα recruitment (Figure 5E, lane 10).

## Discussion

In vertebrates, there are four homologs of Cornichon: CNIH-1, 2, 3, & 4. CNIH-2 and CNIH-3 was shown to play a role in regulating the subunit composition of AMPA receptors, presumably by regulating trafficking of the receptor components from the ER (Kato *et al*., 2010; Herring *et al*., 2013). CNIH-4 was shown to play a role in the ER export of G protein coupled receptors (GPCRs) in HeLa cells (Sauvageau *et al*., 2014). These observations suggest that Cornichon homologs play an evolutionarily conserved role as cargo receptors in ER export. However, overexpression of CNIH in HeLa cells increases the retention of proTGFα in the ER (Castro *et al*., 2007). This may be explained by our observation that overexpression of CNIH results in its ER retention, which in turn traps proTGFα in the ER (Figure S1). By analogy, this may also explain the observation that CNIH-4 overexpression in HeLa cells similarly results in the ER retention of GPCRs (Sauvageau *et al*., 2014).

We show that the efficient recruitment and packaging of proTGFα requires the cooperation of CNIH with one or more cytosolic factors. The yeast homolog, Erv14p, is required for the ER export of the transmembrane proteinYor1p, even though Yor1p interacts directly with Sec24p. The requirement of dual ER export signals is known for other Erv14p client proteins as well, suggesting that this phenomenon was not restricted to Yor1p (Pagant *et al*., 2015). Our observations suggest that this may also be the case for CNIH client proteins in mammals. This may be due to the weak and transient nature of interactions between cargo and coats, which requires multiple sites of interaction to stabilize the pre-budding complex for efficient incorporation into COPII vesicles.

Although a defect in pro/HA-TGFα recruitment and packaging was observed *in vitro*, a significant defect in pro/HA-TGFα steady state localization or ER export kinetics was not detected in CNIH KO cells (Figure S2). We did find evidence of a change in the cytosolic protein content of CNIH KO cells, which resulted in elevated COPII activity *in vitro* (Figure 4A), which supports the idea that the cells had undergone adaptation in response to a loss of CNIH. However, the cause of this change is unclear. Notably, most known client proteins of Erv14p, the yeast homolog of CNIH, are single-pass transmembrane proteins that ultimately localize to the plasma membrane (Powers and Barlowe, 1998; Herzig *et al*., 2012). These plasma membrane localized proteins possess transmembrane domains that are longer than those of ER or Golgi resident proteins (Sharpe *et al*., 2010). Thus it has been proposed that Erv14p may also act as a chaperone in the ER to shield the hydrophobic transmembrane domains of its client proteins (D’Arcangelo *et al*., 2013). Indeed, deletion of the *ERV14* gene was found to enhance the unfolded protein response (UPR) in yeast, which leads to upregulation of many genes involved in the ER-to-Golgi trafficking pathway, including *SEC12, SEC13*, and *SEC24*(Travers *et al*., 2000; Jonikas *et al*., 2009). It is possible that CNIH plays a similar role in mammalian cells, and the loss of CNIH may lead to a moderate UPR in mammalian cells that enhances COPII activity.

The moderate UPR triggered by loss of CNIH may also potentially explain the unexpected impairment of ERGIC53 export (Figure 2D). ERGIC53 oligomerization is required for its efficient export from the ER (Nufer *et al*., 2003). Whereas the COPII interaction motif resides in the cytoplasmic C-terminus of ERGIC53 (Kappeler *et al*., 1997; Wendeler *et al*., 2007), the luminal domain plays a crucial role in mediating oligomerization (Neve *et al*., 2005). Thus it is possible that CNIH KO resulted in changes in the ER lumen that affected ERGIC53 oligomerization, resulting in the diminished budding efficiency observed.

Notably, the V160G mutant form of proTGFα, although packaged inefficiently into COPII vesicles, is recruited normally to the pre-budding complex when CNIH and the cytosolic factor(s) are present. This shows that efficient recruitment of cargo to the pre-budding complex does not guarantee efficient cargo packaging into COPII vesicles, suggesting that cargo recruitment and cargo packaging are distinct processes.

Also noteworthy is the lack of correlation between cargo recruitment and budding efficiency. In the *in vitro* budding reactions, ERGIC53 was very efficiently packaged into COPII vesicles. However, it was undetectable in the COPII binding assays (data not shown). In contrast, Sec22 was also very efficiently packaged into COPII vesicles, while detected at reasonable levels in the COPII binding assays (Figure 5E). This may be explained by the stable association of Sec22 with Sec23/24 upon formation of a fusogenic SNARE complex with Bos1 & Sed5, as shown with yeast proteins (Sato and Nakano, 2005). The apparently different binding affinities of two efficiently packaged cargo proteins to the pre-budding complex again supports the notion that cargo recruitment and cargo packaging are two distinct events in the process of COPII vesicle formation.

Pre-budding complexes are maintained in a highly dynamic manner. Interaction between Sec24 and cargo persists even after Sar1 hydrolyzes its bound GTP, whereas Sec12 activity dynamically maintains the interaction between Sec24 and cargo, presumably by reloading Sar1 with GTP (Sato and Nakano, 2005). This suggests that multiple rounds of GTP hydrolysis by Sar1 may be required for productive packaging of cargo into COPII vesicles. The established methods used to isolate pre-budding complexes utilize either non-hydrolyzable GTP analogs or GTPase defective Sar1 mutants, and thus are unable to capture the dynamic nature of this assembly intermediate and may explain the extremely low efficiency of its recovery (Aridor *et al*., 1998; Kuehn *et al*., 1998).

The selection of appropriate cargo proteins to include into nascent vesicles is a critical function of the COPII coat, in addition to its role in vesicle formation. This is the fundamental mechanism to ensure proper functioning of the eukaryotic endomembrane system, and by extension, to ensure cell viability. During the course of evolution, novel mechanisms may have emerged to cope with the expanding repertoire of cargo proteins that pass through the endomembrane system. New methods are needed to capture the transient and dynamic nature of cargo recruitment and packaging. The identification and characterization of the auxiliary cytosolic factor(s) required for efficient COPII packaging of proTGFα will also provide important information to help elucidate the molecular mechanisms for cargo recognition and packaging in mammalian cells.

## Materials and Methods

### Antibodies

Anti-Sec13 and anti-Sec22 sera were prepared as previously described (Merte *et al*., 2010). Anti-Sec31A was from BD Transduction Laboratories (San Jose, CA) (612350). Anti-Sec23A, anti-Sar1A/B and anti-ERGIC53 sera were prepared as previously described (Fromme *et al*., 2007). Anti-ribophorin I serum was provided by Peter Walter (UCSF). Rabbit anti-HA monoclonal antibody was from Cell Signaling (Danvers, MA) (C29F4). Anti-CNIH was from Sigma (St. Louis, MO) (SAB1304796). Anti-mouse and anti-rabbit HRP conjugates were from GE Healthcare (Buckinghamshire, UK) (NXA931 and NA934V, respectively).

### Preparation of rat liver cytosol

Where indicated, rat livers were preserved by flash freezing with liquid N_2_ and stored at −80°C. Frozen livers were then thawed in 400ml PBS at 4°C. Fresh livers were used unless otherwise indicated. Rat liver cytosol was prepared in Buffer E as described (Kim *et al*., 2005), typically at a protein concentration of 30-40 mg ml^−1^.

### Preparation of HeLa cytosol

Wild type or CNIH knockout HeLa cells were grown in ten 150mm plates to 90% confluency, washed with PBS, and collected using a cell scraper into 10ml B88 buffer containing 1x protease inhibitors (PI). Cells were rendered permeable by exposure to 80μg ml^−1^ digitonin in B88 with PI for 30min at 4°C with gentle rocking. Bio-Beads SM-2 (2g) (Bio-Rad) resin were hydrated and washed with 35ml B88 buffer. Permeabilized cells were centrifuged at 300 *g* for 5min at 4°C, and the supernatant (10ml) was collected and incubated with Bio-Beads overnight at 4°C with gentle rocking to remove cell debris and digitonin. Bio-Beads were subsequently removed by centrifugation at 300 *g* for 5min at 4°C. The resulting supernatant was further centrifuged at 135,000 *g* for 30min at 4°C to remove remaining insoluble material or protein aggregates. The supernatant was then transferred to Amicon tubes and concentrated to a final volume of ~500μl Protein concentration was then measured and the final cytosol was distributed into aliquots, flash frozen, and stored at −80°C.

### Preparation of permeabilized cells

Procedure for preparing permeabilized cells and subsequent *in vitro* translation was performed as previously described (Merte *et al*., 2010), with slight modifications. In detail, HeLa cells were grown in a 100mm dish to 90% confluency, washed with 5ml PBS, trypsinized at RT for 5min, and washed again in 6ml KHM buffer containing 10μg ml^−1^ soybean trypsin inhibitor. Cells were then permeabilized in 40μg ml^−1^ digitonin for 5min in 6ml ice cold KHM buffer. Permeabilized cells were then washed once with B88-LiCl buffer, once in B88 buffer, and finally resuspended in B88 buffer such that the optical density (at 600nm) of membranes in B88 buffer was 0.750. The resuspension was aliquoted and stored at −80°C.

For use in *in vitro* translation, semi-intact cells were washed once in KHM buffer after permeabilization with digitonin and resuspended in 100 μl KHM buffer.

### *In vitro* translation

*In vitro* translation was performed as previously described (Merte *et al*., 2010), with slight modifications. In detail, 1mM CaCl_2_ and 10μg ml^−1^ micrococcal nuclease were added to 100μl of permeabilized cells prepared as described above to remove endogenous RNA. After incubation at room temperature for 12min, 4mM EGTA was then added to terminate the reaction. Membranes were then centrifuged at 10,000 *g* at 4°C for 15s, and resuspended in KHM buffer such that the optical density (at 600nm) of 5 μl membranes in 500μl KHM buffer was 0.060. These membranes were added to *in vitro* translation reactions containing rabbit reticulocyte lysate (Promega, Flexi), amino acids, KCl, and mRNA encoding HA-Vangl2, which was incubated at 30°C for 60min (Merte *et al*., 2010). Donor membranes were centrifuged at 2,700 *g* at 4°C for 3min, and resuspended with 1ml B88-LiCl buffer. Donor membranes were then washed twice with B88 buffer, and resuspended in 20μl B88 buffer per reaction.

### *In vitro* COPII vesicle budding assay

Vesicle formation and purification was performed as described previously (Kim *et al*., 2005), with slight modifications. In detail, donor membranes were incubated where indicated with rat liver cytosol (2 mg ml^−1^), recombinant COPII proteins (10 ng μl^−1^ Sar1A, 10 ng μl^−1^ Sec23A/24D, 10 ng μl^−1^ Sec13/31A), ATP regeneration system, 0.3mM GTP, and Sar1A H79G (10 ng μl^−1^) in a final volume of 100μl. The reaction was carried out at 30°C for 1h, and terminated by incubation on ice for 5min. Donor membranes were removed by centrifugation at 14,000 *g* for 12min at 4°C. The resulting supernatant was centrifuged at 115,000 *g* for 25min at 4°C to collect COPII vesicles. Vesicle pellets were then resuspended in 15μl of Buffer C and Laemmli sample buffer and heated at 55°C for 15min. Vesicle samples and original donor membrane samples were resolved by SDS-PAGE, transferred on to PVDF membranes, and subjected to immunoblotting to detect various ER resident proteins and COPII cargo proteins.

### COPII recruitment assay

The membrane recruitment of COPII components using recombinant Sar1A-GTP-restricted mutant was performed as previously described (Aridor *et al*., 1995), with modifications. In detail, donor membranes were mixed where indicated with rat liver cytosol (2 mg μl^−1^), recombinant COPII components (20 ng μl^−1^ Sec23A/24D), ATP regeneration system, 1mM GTP, and GST-Sar1A-GTP-restricted mutant protein (30 ng μl^−1^) in a total volume of 100μl. The mixture was incubated at 30°C for 30min, after which the reaction was terminated by transfer to ice. Membranes were collected by centrifugation at 20,000 *g* for 10min at 4°C. The supernatant was discarded, and membranes were dissolved by incubation with 1ml of solubilization buffer (20mM HEPES, pH 7.2, 1mM magnesium acetate, 1% digitonin) on ice for 30min, with occasional mixing. Insoluble material was removed by centrifugation at 163,000 *g* for 30min at 4°C. The resulting supernatant was then incubated with glutathione agarose beads (Pierce, Waltham, MA) at 4°C for 30min with rotational rocking. Subsequently, the beads were collected by centrifugation (200 *g*) and washed three times with 200μl solubilization buffer. Beads were eluted by heating in Laemmli sample buffer at 55°C for 15min. Samples were resolved by SDS-PAGE, transferred on to PVDF membranes, and subjected to immunoblotting to detect various ER resident proteins, COPII cargo proteins, and Sec23A.

### Purification of Recombinant Proteins

Purification of human Sar1A WT & H79G was performed as described (Kim *et al*., 2005). Purification of human GST-Sar1A-GTP-restricted form was performed as described (Kim *et al*., 2005), with modification. Briefly, instead of thrombin cleavage, the GST-tagged recombinant protein was directly eluted with 5mM glutathione in PBS buffer, dialyzed overnight in HKM-G buffer, distributed into aliquots, flash frozen in liquid N_2_, and stored at −80°C. Purification of human FLAG-Sec23A/His-Sec24D was performed according to previously described procedures (Kim *et al*., 2005).

### Immunoprecipitation

HeLa cells grown in 10cm plates were transiently transfected with 2°g of plasmids encoding pro/HA-TGFα wild type or V160G mutant, or HA-Vangl2, using Lipofectamine 2000 according to the manufacturer’s recommended protocol. Twenty-four hours later, cells were washed twice in PBS. Cells were then solubilized in 1ml of IP lysis buffer (20mM Tris-HCl, pH 7.4, 150mM NaCl, 1mM EDTA, 0.5% Triton X-100, 1x Protease Inhibitors) on ice for 15min. Cell lysates were cleared by centrifuging at 20,817 *g* for 15min at 4°C. An aliquot (10μ1) of the supernatant fraction was reserved in a separate tube and mixed with Laemmli sample buffer. An aliquot (30μ1) of 50% monoclonal anti-HA-agarose (Sigma-Aldrich) slurry was washed three times with IP lysis buffer and added to cleared cell lysates and incubated at 4°C for 1h. The monoclonal anti-HA-agarose beads were centrifuged at 200 *g* and washed three times with IP lysis buffer. The beads were then resuspended in 15μl 1 × Laemmli sample buffer, heated to 55°C for 15min, and samples were resolved by SDS-PAGE, transferred onto PVDF membranes and subjected to immunoblotting.

## Acknowledgements

We thank Lixon Wang, Lazar Dimitrov, and Kartoosh Heydari for their assistance in preparing CNIH KO HeLa cells. We thank Ann Fisher at the Berkeley Tissue Culture Facility for assistance with cell culture. The plasmid encoding CNIH-HA was a generous gift from Carolina Castro and Rik Derynck.

## Supplementary Methods

### TMR-Star Pulse-Chase

Cells were transfected with a plasmid encoding pro/HA-SNAP-TGFα. At 24h post transfection, normal growth media was exchanged with growth medium containing 2μM of SNAP-Cell® Block (NEB, S9106S) for 30min, to block the SNAP activity of all previously synthesized pro/HA-SNAP-TGFα. Afterwards, cells were washed 5x with growth media and incubated in growth media containing 1.5μM of SNAP-Cell® TMR-Star (NEB, S9105S) to label newly synthesized pro/HA-SNAP-TGFα. After a certain amount of time (as indicated in Figure S2), cells were fixed and prepared for microscopy.

### Immunofluorescence microscopy

Cells were fixed in 3% paraformaldehyde (PFA) for 30min and then washed 5x with 1xPBS, then permeabilized and blocked in blocking buffer (200mM glycine, 0.1% Triton X-100, 2.5% FBS in 1x PBS buffer) for 20min. Cells were then incubated for 30min with rabbit anti-HA antibody (Cell Signaling, C29F4) at 1:200 dilution in blocking buffer, washed, and further incubated with anti-rabbit IgG FITC conjugate (Jackson ImmunoResearch, 711-095-152) diluted 1:250 in blocking buffer for 30min, then washed 5x with 1xPBS. Cells were mounted onto glass slides with ProLong® Gold antifade mountant (ThermoFisher, P36930) and imaged using a Zeiss AxioObserver Z1 fluorescent microscope. Images were captured and merged using Metamorph software from Molecular Devices.

Budding reactions were performed as described in Materials and Methods.

**Figure S1.**
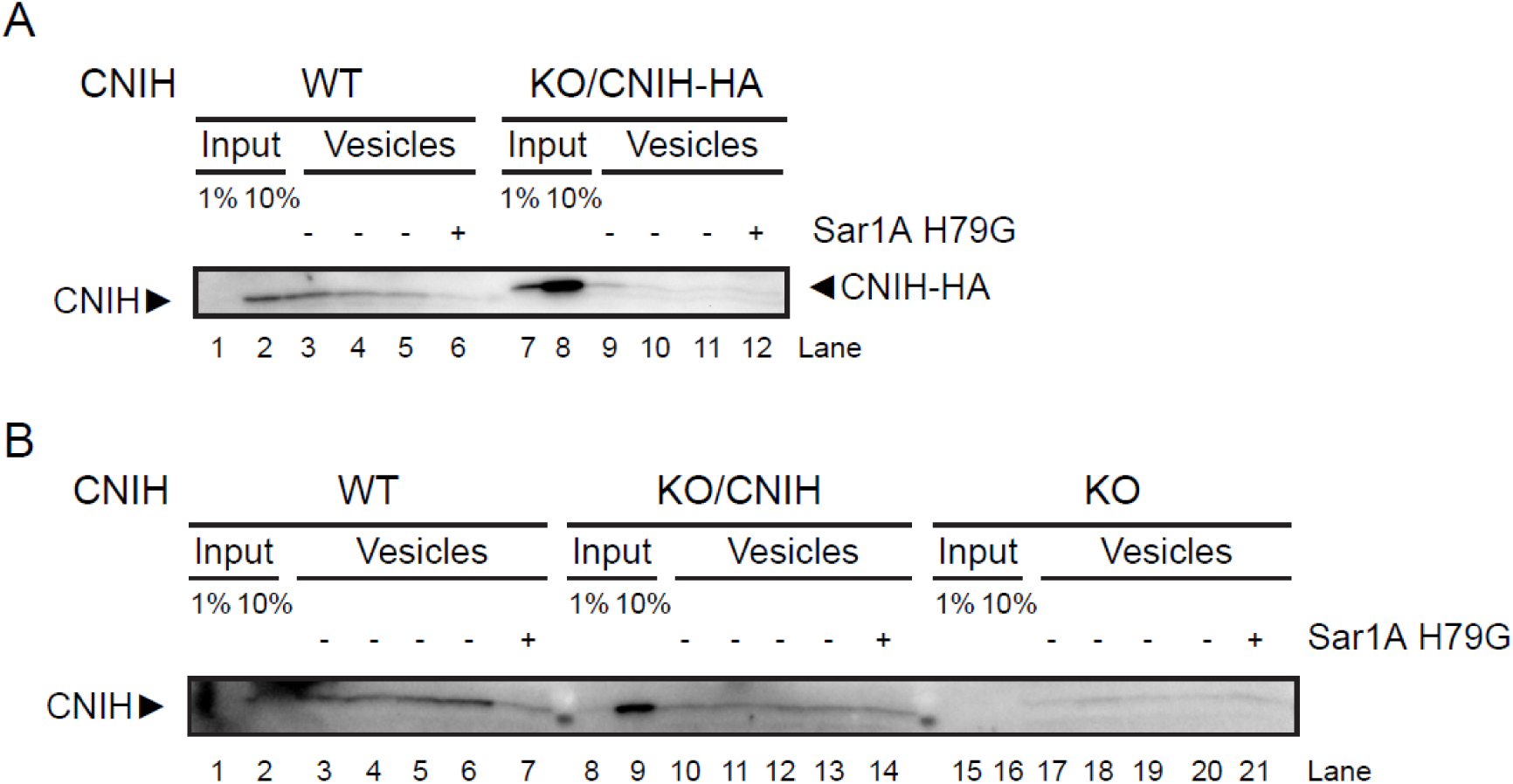
Overexpression of Cornichon-1 results in retention in the ER. (A and B) Immunoblots of in vitro budding reactions. Membranes were prepared as indicated either from wild type (WT) or CNIH KO (KO) HeLa cells. CNIH knockout cells were transiently transfected with plasmids expressing CNIH-HA (KO/CNIH-HA) or CNIH (KO/CNIH), and membranes were harvested 20h post transfection for use in in vitro budding reactions. Overexpression of CNIH-HA resulted in significantly decreased budding efficiency (A). This was not due to addition of the HA-tag as overexpression of CNIH also resulted in very inefficient CNIH budding (B).

**Figure S2.**
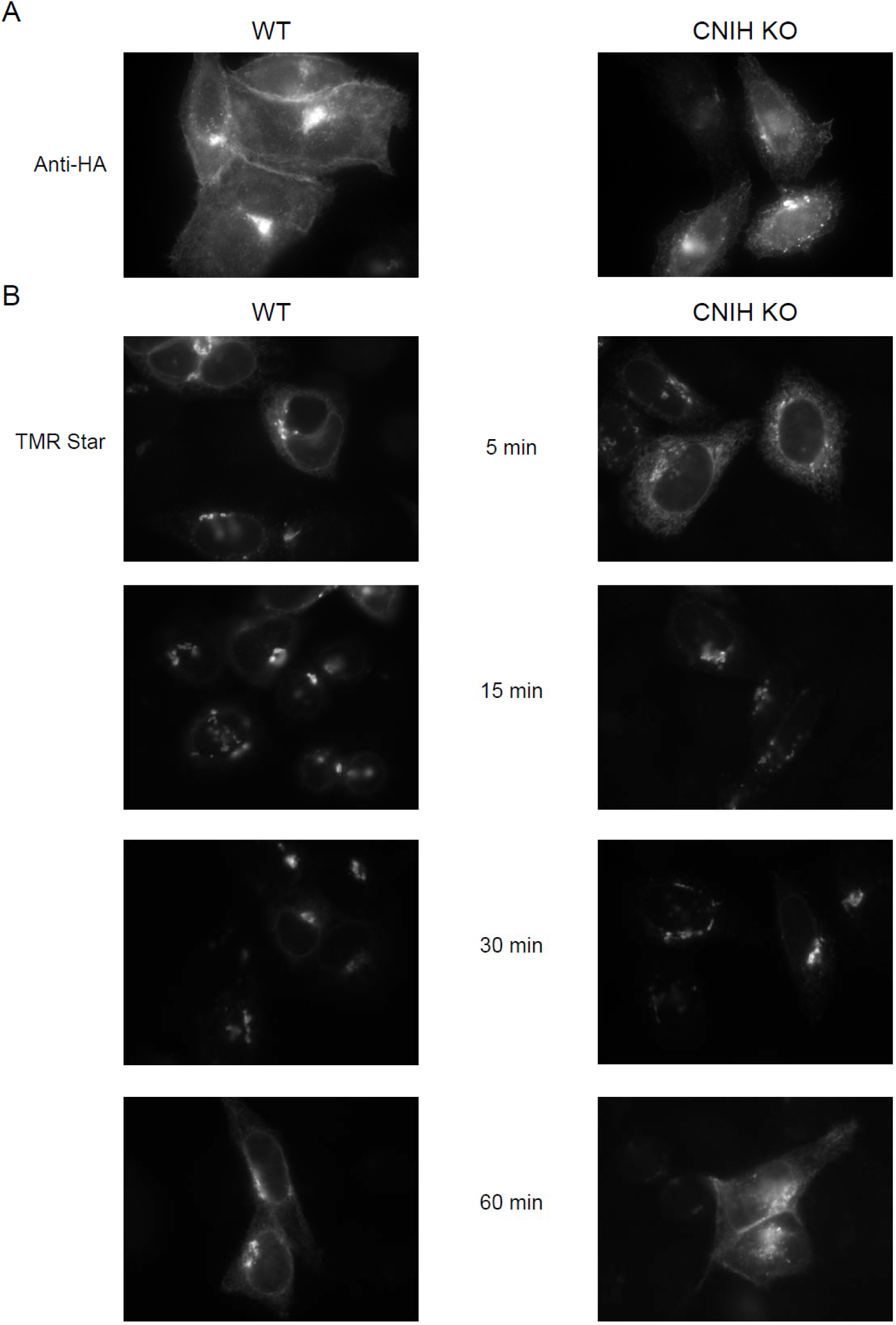
Comparison of pro/HA-SNAP-TGFα localization and trafficking kinetics in WT and CNIH KO HeLa cells. (A) Immunofluorescence localization with anti-HA showing steady state localization of pro/HA-SNAP-TGFα in WT or CNIH KO cells as labeled. The steady state localization of pro/HA-SNAP-TGFα did not show a significant difference. (B) TMR-Star was used to label newly synthesized pro/HA-SNAP-TGFα in WT or CNIH KO cells as indicated. Cells were fixed for microscopy at the indicated time points. At 5min, the majority of newly synthesized pro/HA-SNAP-TGFα was localized in the ER, with some punctate distribution, possibly representing the ERGIC. At 15min and 30min the majority of TMR-Star signal was found in perinuclear punctate structures resembling the Golgi apparatus. At 60min, TMR-Star signal became visible at the plasma membrane, although significant signal was also observed in ER and Golgi-like punctate structures, likely representing newly synthesized pro/HA-SNAP-TGFα in earlier steps of maturation. The differences between pro/HA-SNAP-TGFα localization in WT and CNIH KO cells were subtle.

